# Systematic Evaluation of Cell Type Deconvolution Methods for Plasma Cell-free DNA

**DOI:** 10.1101/2024.03.25.586507

**Authors:** Tongyue Sun, Jinqi Yuan, Yacheng Zhu, Shen Yang, Junpeng Zhou, Xinzhou Ge, Susu Qu, Wei Li, Jingyi Jessica Li, Yumei Li

## Abstract

Plasma cell-free DNA (cfDNA) is derived from cellular death in various tissues. Investigating the origin of cfDNA through tissue/cell type deconvolution allows us to detect changes in tissue homeostasis that occur during disease progression or in response to treatment. Consequently, cfDNA has emerged as a valuable noninvasive biomarker for disease detection and treatment monitoring. Although there are numerous methylation-based methods of cfDNA cell type deconvolution available, a comprehensive and systematic evaluation of these methods has yet to be conducted. In this study, we thoroughly benchmarked five previously published methods: MethAtlas, cfNOMe, CelFiE, CelFEER, and UXM. Utilizing deep whole-genome bisulfite sequencing data from 35 human cell types, we generated cfDNA mixtures with known ground truth to assess the deconvolution performance under various scenarios. Our findings indicate that different factors, including sequencing depth, reference marker selection, and reference completeness, influence cell type deconvolution performance. Notably, omitting cell types present in a mixture from the reference leads to suboptimal results. Despite each method exhibited distinct performances under various scenarios, CelFEER and UXM exhibit overall superior performance compared to the others. In summary, we comprehensively evaluated factors influencing methylation-based cfDNA cell type deconvolution and proposed general guidelines to maximize the performance.

## Introduction

Plasma cell-free DNA (cfDNA) predominantly consists of double-stranded DNA fragments released by dying cells from various tissues^1–3^. In healthy individuals, plasma cfDNA is primarily derived from apoptosis of normal hematopoietic cells, with minimal contributions from other cell types^4–6^. However, the progression and treatment of numerous diseases, including cancers^7^, can impact the distribution of cell-type-derived cfDNA fractions. Consequently, abnormal distributions can serve as indicators of altered tissue homeostasis resulting from disease development or treatment. As a result, cfDNA has emerged as a valuable noninvasive biomarker for disease detection and treatment monitoring^8–10^. Understanding the cell type origin of cfDNA is crucial in elucidating abnormal distribution in patients, contributing to the discovery of noninvasive biomarkers and enhancing our understanding of disease development. Cell type deconvolution is a type of analysis focused on offering a comprehensive composition of different cell types present in cfDNA.

While each cell possesses nearly identical DNA sequence, DNA methylation signatures exhibit cell type specificity, providing a means to trace back to the cell type origin of cfDNA^6,11,12^. The most prevalent form of DNA methylation is 5-methylcytosine (5mC), which predominantly occurs at CpG sites in humans^13^. Several studies have introduced methylation-based deconvolution methods to estimate the proportions of various cell types in cfDNA^5,6,14–16^.

MethAtlas^5^ models the plasma cfDNA methylation profile as a linear combination of the methylation profiles from cell types in the reference atlas. Subsequently, the relative contribution of each cell type to cfDNA is determined using non-negative least squares linear regression (NNLS). CfNOMe^15^ follows a similar approach to MethAtlas but utilizes linear least squares minimization to optimize each cell type’s relative contribution. CelFiE^14^ estimates the contribution of different cell types to the cfDNA via an expectation-maximization optimization algorithm. Notably, CelFiE can also estimate contributions from multiple unknown cell types not available in the reference atlas. CelFEER^16^, an adapted method from CelFiE, utilizes average methylation values within individual reads instead of the methylation ratio of CpG sites to differentiate cell types. UXM is a fragment-level deconvolution algorithm using the percentage of unmethylated fragments for methylation quantification of reference markers.

Nevertheless, a comprehensive assessment of the performance of these methods has not yet been conducted. Additionally, each method necessitates specific preprocessing procedures and relies on distinct reference methylation atlases. The potential impact of these procedures on deconvolution results remains unknown. To address this knowledge gap, we conduct a thorough comparison and evaluation of five methylation-based cfDNA cell type deconvolution methods with publicly available and executable codes (MethAtlas, cfNOMe, CelFiE, CelFEER, and UXM) (**Fig. 1, Supplementary Table 1**). Our evaluation encompasses their performance in terms of reference marker selection outcomes and deconvolution accuracy under various scenarios, utilizing in silico mixtures of cell types’ whole-genome methylation sequencing (WGBS) data with known ground truth. Based on the analytical findings, we finally summarize the strengths and weaknesses of different deconvolution methods, offering a general guideline for users.

**Fig. 1.**
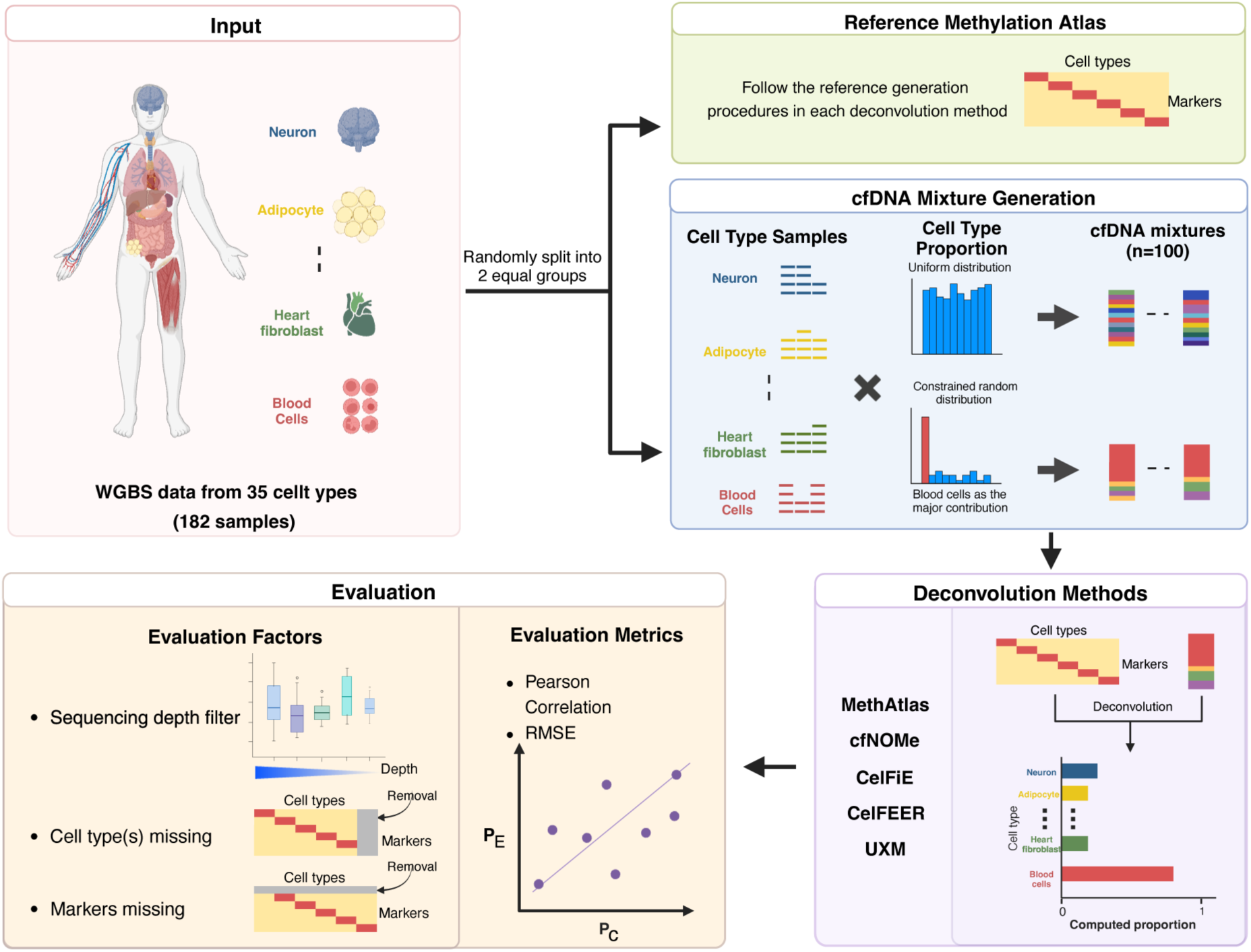
Schematic overview of the benchmarking study. In this benchmarking study, WGBS data from 182 samples representing 35 cell types is initially randomly divided into two equal groups. One group is designated for the creation of the reference methylation atlas, while the other is utilized for generating in silico cfDNA mixtures. Known cell type proportions are then generated using either a uniform distribution or a constrained random distribution with blood cells as the primary contributors. Subsequently, the deconvolution performance is rigorously assessed under various influencing factors, including sequencing depth filter thresholds, cell types missing, and markers missing in the reference methylation atlas. Two evaluation metrics, Pearson correlation and root-mean-square error (RMSE), are employed to scrutinize the accuracy of computed proportions (𝑃_𝑐_) against expected proportions (𝑃_𝐸_).

## Results

### Design of the benchmarking study

The benchmarking analysis utilized a recently published DNA methylation atlas of normal human cell types based on deep WGBS^6^. As shown in **Fig. 1**, from this atlas, we selected 35 cell types, totaling 182 samples, each with at least two biological replicates. To ensure independent reference and testing datasets, we randomly divided all samples into two equal groups: one for generating the reference methylation atlas and the other for creating in silico cfDNA mixtures.

The reference methylation atlas was generated following the procedures specified by each deconvolution method. To assess the impact of cell type proportion distribution, we created two cfDNA mixture datasets (n = 100 for each): 1) a uniform-distribution dataset, with cell type proportions determined by a uniform distribution (**Supplementary Table 2**); and 2) a constrained-random-distribution dataset, emphasizing blood cells as the major contributors to mimic the cfDNA biological scenario in healthy individuals (**Supplementary Table 3, Fig. 1**).

We assessed the performance of different deconvolution methods using Pearson correlation and root-mean-square error (RMSE) values between each method’s computed cell type proportions and the known compositions of cfDNA mixtures. Furthermore, to comprehensively evaluate the influence of various factors on the final result, we scrutinized different sequencing depth filter thresholds, missing cell types, and markers absent during reference marker selection (**Fig. 1**).

### Evaluation of deconvolution accuracy and computing resource requirements

To begin with, we assessed the overall performance of each deconvolution method using their default settings. In the case of the uniform-distribution dataset, CelFEER outperformed others, achieving a median Pearson correlation of 0.82 and an exceptionally low RMSE of 0.0097 (**Fig. 2A and C**). Conversely, CelFiE exhibited the least favorable performance, with median RMSE value nearly double that of CelFEER (**Fig. 2A and C**). Notably, CelFEER demonstrated lower standard deviations for both Pearson correlation and RMSE values across the 100 cfDNA mixtures, indicating its superior stability compared to other methods (**Supplementary Table 4**). For the constrained-random-distribution dataset, overall performance improved compared to the uniform-distribution dataset, with UXM leading the way, which outperformed the other four methods (**Fig. 2B and D**).

**Fig. 2.**
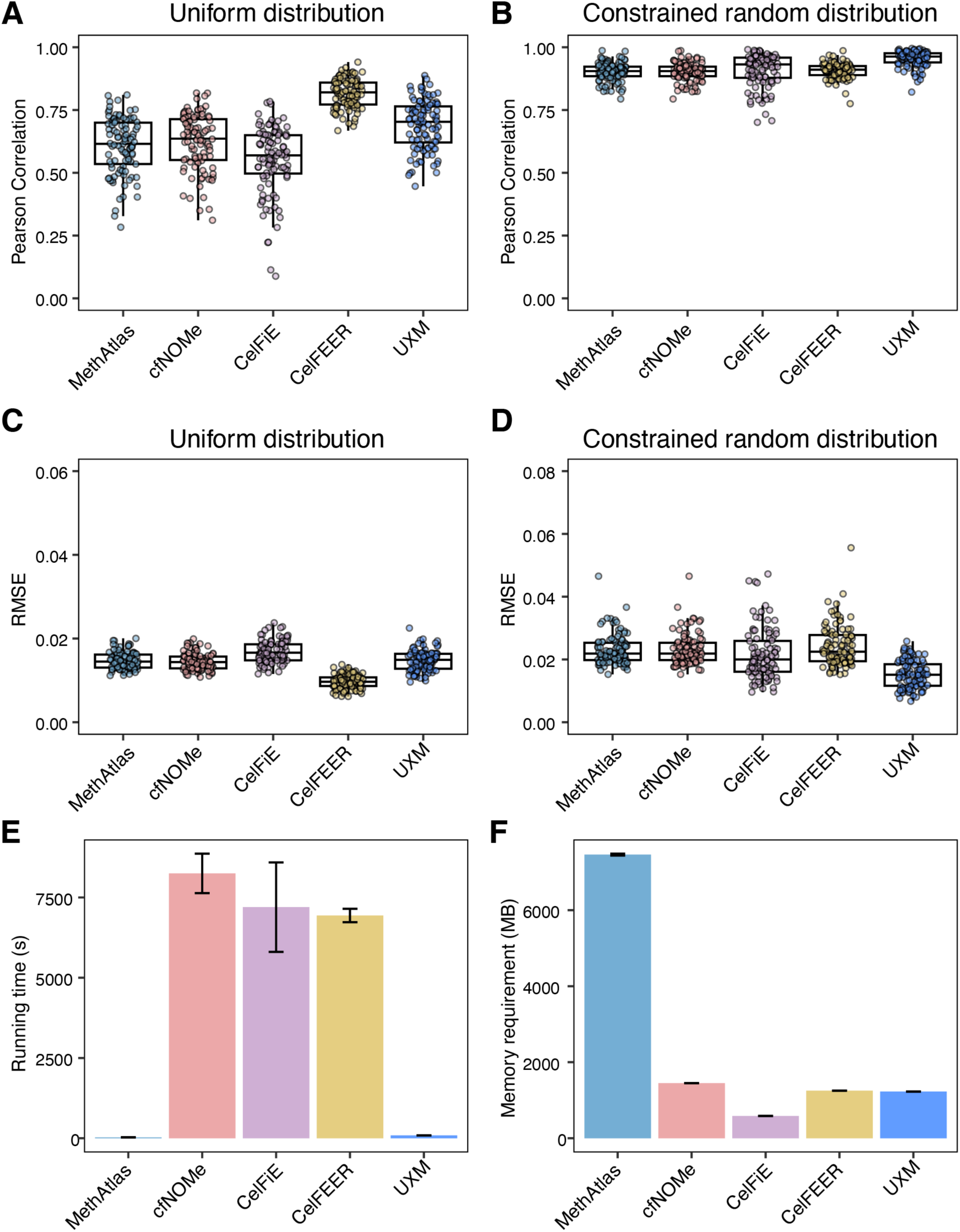
Deconvolution results under the default setting of evaluated methods. **(A-B)** Pearson correlation between the known proportions of the reference cell types in the 100 cfDNA mixtures and the computed proportions from various deconvolution methods. The known proportions are generated under uniform **(A)** and constrained random distributions **(B)**. **(C-D)** RMSE values between the known proportions of the reference cell types in the 100 cfDNA mixtures and the predicted proportions from various deconvolution methods. The known proportions are generated under uniform **(C)** and constrained random distributions **(D)**. RMSE, root mean squared error. **(E)** Running time for various deconvolution methods. **(F)** Memory requirements for various deconvolution methods.

Turning to computing resource requirements, UXM and MethAtlas significantly outpaced the other three methods in terms of running time for analyzing a set of 100 cfDNA mixtures (**Fig. 2E**). Unsurprisingly, MethAtlas, with the most straightforward optimization problem, was the fastest, completing the analysis of 100 samples in under 30 seconds. Conversely, cfNOMe was the slowest, requiring an average of 1.4 minutes to analyze one sample. In terms of memory usage, MethAtlas had the highest RAM requirements, followed by cfNOMe (**Fig. 2F**). CelFiE, on the other hand, exhibited minimal memory usage as low as 585 MB, making it suitable for execution on a personal computer (**Fig. 2F**).

### Evaluation of reference marker selection across methods

Selecting reference markers for each cell type plays a pivotal role in shaping the final deconvolution results. As depicted in **Fig. 3A**, substantial differences in selected markers were evident among the methods, with CelFiE having an order of magnitude more markers than those used in UXM. This discrepancy indicates that different methods may capture distinct cell-type-specific methylation features.

**Fig. 3.**
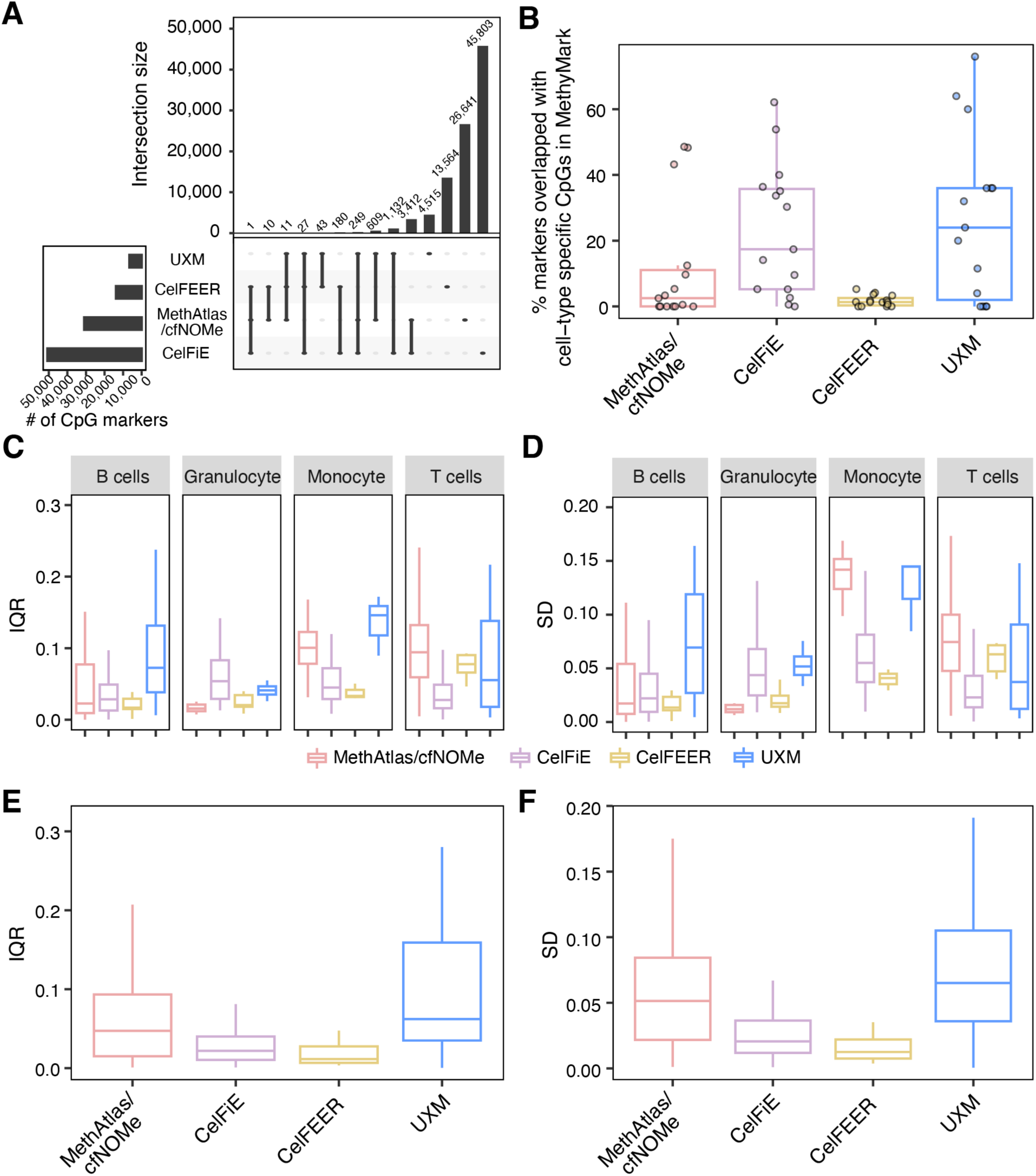
Evaluation of reference marker selection in each deconvolution method. **(A)** Upset plots illustrate the intersections of selected markers from various methods. **(B)** Overlaps between the selected markers from different methods and cell-type-specific CpGs from the MethyMark database. The proportions of overlaps across 15 cell types, as cataloged in MethyMark, are detailed. CfNOMe used the same set of reference markers as MethAtlas. **(C-D)** The variability of the selected markers across healthy individuals was measured by Inter-Quartile Range (IQR) **(C)** and Standard Deviations (SD) **(D)**. IQR and SD data are sourced from the ImmuMethy database, focusing on four immune cell types. **(E-F)** The variability of the selected markers across diverse healthy states was measured by IQR **(E)** and SD **(F)**. Data for IQR and SD are extracted from the ImmuMethy database.

To evaluate the quality of reference marker selection, we hypothesized that ideal reference markers should align with cell-type-specific CpGs, exhibiting significant methylation pattern differences between cell types. Therefore, we primarily assessed reference marker selection results by comparing them with the cell-type-specific methylation marks in the MethyMark database^17^. MethyMark integrated 50 methylomes across 42 human tissues/cell types, 15 of which were also included in the reference methylation atlas of our study (**Supplementary Table 5**). For each of the 15 overlapping cell types, we examined the concurrence between reference markers identified by each deconvolution method and cell-type-specific markers in MethyMark. Notably, UXM identified a higher proportion of reference markers that also corresponded to cell-type-specific markers in MethyMark (**Fig. 3B**), suggesting its potential preference for capturing cell specificity in methylation levels.

Moreover, an optimal reference methylation atlas should include markers that remain stable across individuals and diverse conditions. Consequently, we assessed reference marker selection results by examining marker variability based on data in ImmuMethy^18^, which collected DNA methylation data for immune cells in different conditions. Variability was quantitatively measured using the inter-quartile range (IQR) and standard deviations (SD). Among the four immune cell types (B cell, granulocyte, monocyte, and T cell) included in the reference methylation atlas of our study and also collected in ImmuMethy with more than three conditions, CelFEER exhibited overall lower variability across individuals in each condition, as evidenced by both IQR and SD metrics (**Fig. 3C, D and Supplementary Figs. 1-4**). CelFEER also demonstrated overall lower variability across different conditions (**Fig. 3E** and **F**). These lower variabilities may contribute, at least in part, to CelFEER’s superior performance. However, UXM, which efficiently captured a significant proportion of cell-type-specific markers in MethyMark (**Fig. 3B**) and demonstrated relatively superior deconvolution performance (**Fig. 2B** and **D**), displayed the greatest variability across diverse conditions (**Fig. 3E** and **F**). In conclusion, while the stability of the reference methylation markers could influence the performance of deconvolution methods, it is not the sole determining factor.

### Impact of sequencing depth filter threshold on deconvolution results

The sequencing depth filter threshold utilized during the identification of reference methylation markers can significantly affect the quality and total number of the final selected markers.

Therefore, we sought to assess whether varying sequencing depth filter thresholds impacted deconvolution outcomes. Starting from the default sequencing depth filter threshold (15X) for most deconvolution methods, we explored different threshold settings up to 100X. Subsequently, we executed the deconvolution pipelines under each sequencing depth threshold, using both the uniform-distribution and constrained-random-distribution datasets.

Notably, UXM exhibited the most stable results for both datasets (**Fig. 4** and **Supplementary Fig. 5**), potentially benefiting from its reference marker selection procedure, which involved selecting a fixed number of markers after the sequencing depth filter. Conversely, for MethAtlas and cfNOMe, higher sequencing depth thresholds were associated with poorer performance (**Fig. 4** and **Supplementary Fig. 5**), indicating the impact of a decreased number of selected markers as the sequencing depth filter threshold increased. CelFiE and CelFEER showed slightly improved performance with increasing filter thresholds, suggesting the influence of reference marker quality for these two methods (**Fig. 4** and **Supplementary Fig. 5**).

**Fig. 4.**
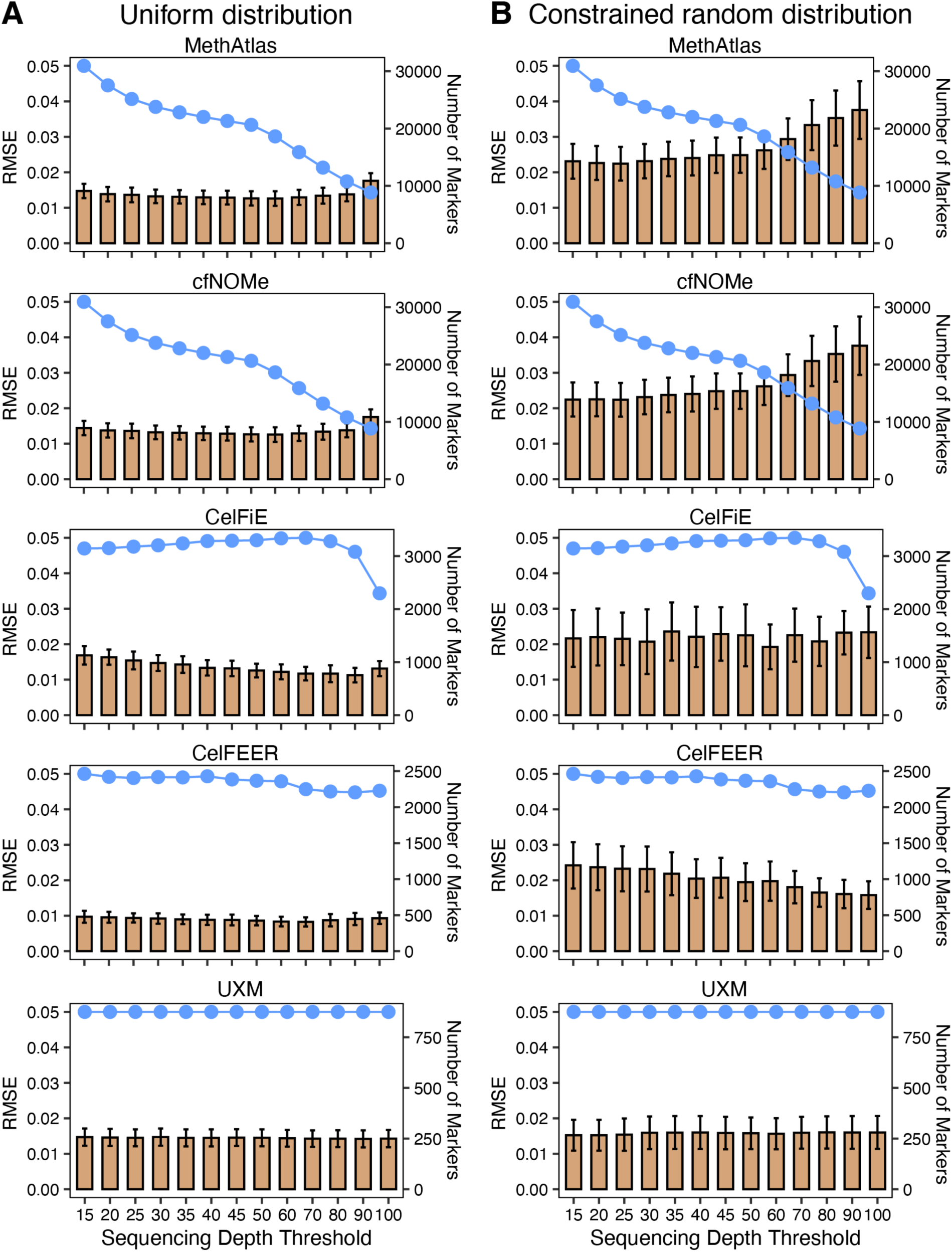
Impact of sequencing depth filter threshold on deconvolution results. **(A-B)** RMSE values between the known and predicted proportions from different deconvolution methods, considering various sequencing depth thresholds for marker selection. The known proportions are generated under uniform **(A)** and constrained random distributions **(B)**. The line plot depicts the number of selected markers.

### Influence of reference atlas incompleteness

Ideally, a comprehensive reference atlas should encompass all informative methylation markers and include as many cell types as possible. In practical terms, however, the presence of missing reference markers or cell types is inevitable due to the constraints imposed by the availability of high-coverage sequencing data. Thus, we conducted an assessment to determine whether the incompleteness of the reference methylation atlas impacted deconvolution results. Firstly, we executed the five deconvolution methods using reference atlases with varying proportions of markers (10%, 20%, 30%, 40%, and 50%) removed from the full atlas. Under both the uniform-distribution and constrained-random-distribution datasets, UXM exhibited diminishing performance with increasing proportions of missing markers (**Fig. 5A** and **B**, **Supplementary Fig. 6**). CelFEER and CelFiE displayed considerable variances in the uniform-distribution dataset but demonstrated stable performance under the constrained-random-distribution dataset, suggesting their instability. In contrast, MethAtlas and cfNOMe showed consistent performance under different datasets and proportions of missing markers (**Fig. 5A** and **B**, **Supplementary Fig. 6**).

**Fig. 5.**
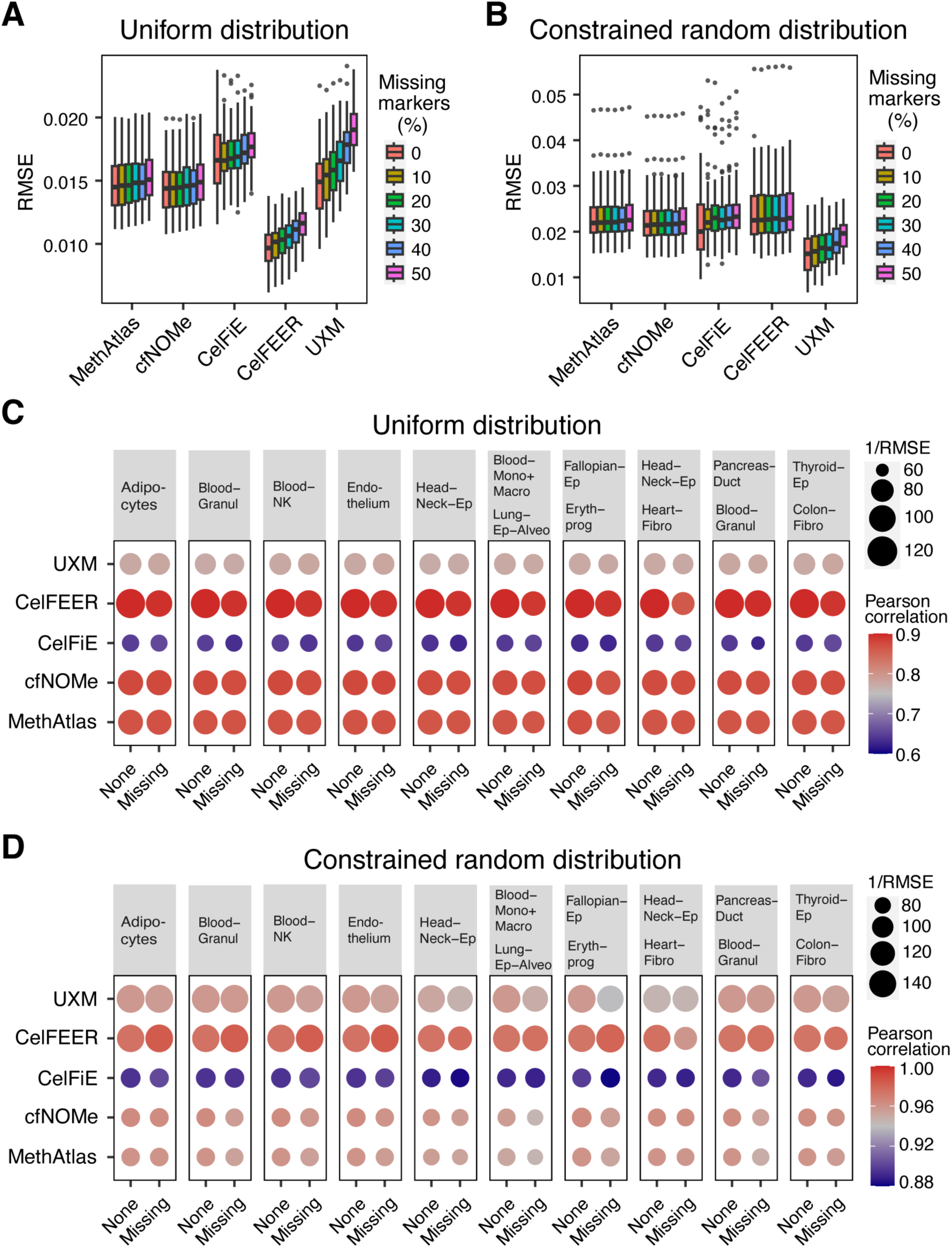
Influence of reference marker incompleteness on deconvolution results. **(A-B)** RMSE values between the known and predicted proportions from various deconvolution methods. Different proportions of markers are intentionally omitted in the reference. The known proportions are generated under uniform **(A)** and constrained random distributions **(B)**. **(C)** Pearson correlation and RMSE values between known proportions under uniform distribution and the predicted proportions from different deconvolution methods with no, one, or two cell types intentionally missing in the reference. None represents no cell type missing scenario. Missing represents one or two cell type missing scenarios. **(D)** Pearson correlation and RMSE values between the known proportions under constrained random distribution and the predicted proportions from different deconvolution methods with no, one, or two cell types intentionally missing in the reference. None represents no cell type missing scenario. Missing represents one or two cell type missing scenarios.

Subsequently, we examined the influence of missing cell types by removing one or two cell types from the reference atlas while keeping the in-silico cfDNA mixtures unchanged. To ensure an equitable comparison, we exclusively calculated the Pearson correlation and RMSE for cell types present in both scenarios of no missing cell types and one/two missing cell types. Among the five methods, only CelFiE and CelFEER could estimate the proportions of unknown cell types not available in the reference atlas. However, CelFEER exhibited poor performance when one cell type was missing and even worse when two cell types were missing in the uniform-distribution dataset (**Fig. 5C**). UXM and CelFiE showed decreased performance when two cell types were missing, while the performances of MethAtlas and cfNOMe remained relatively stable under the uniform-distribution dataset (**Fig. 5C**). Nevertheless, for the constrained-random-distribution dataset, the performance of all methods deteriorated when there were missing cell types, particularly when the missing cell types included blood cells (**Fig. 5D**). To evaluate the impact of missing major cell types in the reference atlas, we subsequently removed each of the five blood cell types or one blood cell type coupled with another randomly selected cell type for the constrained-random-distribution dataset. All methods, except CelFEER, exhibited decreased performances compared to the results when there were no missing cell types (**Supplementary Fig. 7**). CelFEER showed slightly better performance in scenarios where one cell type was missing, potentially benefiting from its ability to estimate unknown cell type proportions. In conclusion, the incompleteness of the reference atlas could influence deconvolution results to varying degrees for different methods and different proportions of the missing cell type.

## Discussion

We conducted a comprehensive evaluation of the performance of five methylation-based cfDNA deconvolution methods using in silico cfDNA mixtures and assessed them based on Pearson correlation and RMSE values. Given that cfDNA primarily originates from apoptosis of diverse cell types, we opted to utilize methylation data of cell types for generating the mixtures.

However, previous methylation datasets often cover only a fraction of genomic regions and are typically based on tissues containing a mixture of cells^19–21^. A recent contribution by Loyfer *et al.* provided a human methylome atlas encompassing 39 cell types derived from 205 healthy tissue samples through deep whole-genome bisulfite sequencing^6^. This atlas proves to be a valuable resource for benchmarking cfDNA deconvolution methods. Notably, the inclusion of biological replicates for most cell types in this atlas allows us to differentiate replicates for constructing the reference atlas and generating mixtures, thereby avoiding the potential "double dipping" problem^22^. Leveraging the deep sequencing depth (∼30X) of this methylome atlas, we can also assess the impact of sequencing depth filters and reference marker incompleteness. Based on the comprehensive benchmarking results, we offer a guideline for researchers to choose suitable tools (**Fig. 6**).

**Fig. 6.**
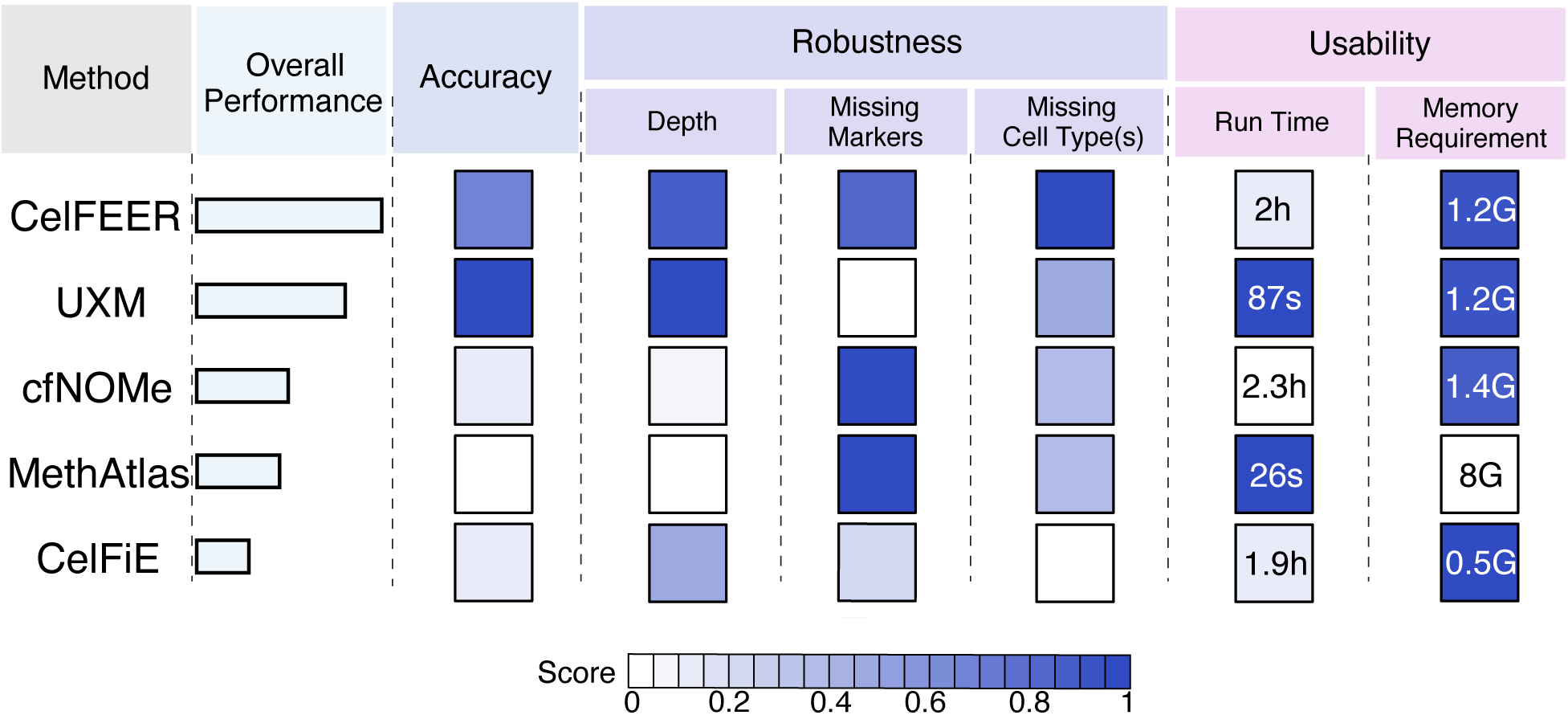
Overall summary of the methods’ performance. A concise overview of the benchmarking results for the five cfDNA deconvolution methods. Refer to the **Methods** section for detailed information on the ranking criteria and methodology.

Our analyses were conducted on two cfDNA mixture datasets to encompass diverse biological scenarios and assess the robustness of these methods. The constrained-random-distribution dataset aimed to mimic the actual cfDNA compositions in healthy individuals^5,6^, while the uniform-distribution dataset was employed to evaluate the methods’ robustness to different cell type compositions. Each method exhibited distinct performances between the two datasets, particularly CelFEER. Additionally, we observed that overall Pearson correlation from the constrained-random-distribution dataset were higher than those from the uniform-distribution dataset, indicating superior performance in the constrained-random-distribution dataset.

However, the RMSE values showed an overall increase from the constrained-random-distribution dataset, signifying poorer performance. This inconsistency underscores the importance of employing multiple evaluation metrics. In practical scenarios, researchers often cannot provide a complete reference atlas containing all the markers and cell types. Therefore, we also evaluated the influence of missing markers and cell types in the reference atlas. While some methods, such as MethAtlas and cfNOMe, demonstrated stable performances under these factors, others proved to be sensitive. Consequently, researchers should carefully consider the completeness of their data when selecting tools for cfDNA deconvolution analysis.

While our benchmarked methods yielded reasonable deconvolution results in most analyses, certain limitations require attention. Firstly, no single method consistently outperformed others across all evaluated factors, implying that the optimization of these methods might prioritize specific considerations. Secondly, the cell-type specific signals are still far from fully captured, which is evident in the limited overlaps between these methods and with the external datasets. Third, none of these methods considers the uncertainty of the deconvolution results which may influence the analytical outcomes. Last but not least, when deploying cfDNA deconvolution tools, the importance of software sustainability and user support must be considered. Notably, some of our benchmarked methods lacked a complete user guide and were less responsive to user inquiries.

In summary, when engaging in cfDNA deconvolution analysis, we recommend users: 1) choose a method stable to sequencing depth, especially with low-coverage data; 2) employ a stringent marker selection strategy to retain the most informative markers; and 3) utilize a comprehensive reference atlas that encompasses all cell types present in the cfDNA.

## Methods

### Data collection and processing

We curated a comprehensive human DNA methylation dataset, including 39 normal human cell types from Gene Expression Omnibus (GEO) with accession number GSE186458^6^. Our selection criteria focused on including cell types having a minimum of two biologically independent replicates of high quality. This rigorous selection process yielded a refined dataset comprising 35 distinct cell types, encompassing a total of 182 samples (**Supplementary Table 6**).

Subsequently, we partitioned all samples into two equal groups. Replicates from the same cell type in each group were merged into unified samples utilizing the wgbstools^6^. This process generated two independent datasets, each serving a specific purpose. One dataset was employed for the creation of in silico cfDNA mixtures, while the other functioned as the reference methylation atlas.

### The generation of in silico cfDNA mixtures and reference methylation atlas

To investigate the impact of cell type proportion distribution, we generated two sets of cfDNA mixture data, each consisting of 100 samples. These sets were based on two distinct distributions: uniform distribution and constrained random distribution. For the uniform-distribution dataset, we employed the *random.uniform* function from the Python package numpy (v 1.24.3)^23^ to create a 35 × 100 matrix, with each column normalized to 1. The constrained-random-distribution dataset was generated through the following steps: 1) selection of the five immune cell types and random inclusion of 1-10 additional cell types; 2) generation of cell type compositions by following Random Distribution using *random* function from Python package random; 3) assignment of the five largest numbers to the five immune cell types, with the remaining numbers allocated to the other cell types; 4) division all numbers by the sum of the random numbers to ensure that cell type fractions summed to 1. The proportion (𝑃_𝑖_) for a cell type 𝑖 was calculated as 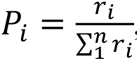, where 𝑟_𝑖_ was the random number generated for the cell type 𝑖. This process was repeated 100 times, resulting in 100 constrained-random-distribution datasets where immune cells were predominant. Subsequently, the mix_pat command in wgbstools was used to generate each cfDNA mixture according to the corresponding proportions. These in silico cfDNA mixtures with known cell type proportions served as the ground truth for subsequent evaluation.

The reference methylation atlas was generated for the five deconvolution methods by following the specific instructions provided by the respective method. In summary:

(1) MethAtlas selected the 100 most hypermethylated and hypomethylated CpGs, along with neighboring CpGs up to 50 bp for each cell type. Pairwise-specific CpGs were then identified through 500 iterations, focusing on pairs of cell types with the smallest Euclidean distance in each iteration.
(2) CfNOMe used the same set of reference markers as MethAtlas.
(3) CelFiE calculated the distance between the methylation ratio of each cell type and the median methylation ratio for all cell types for each CpG. Then, the top 100 CpGs with the largest distance for each cell type, along with the proximal CpGs (±250bp) around these CpGs, were selected to form the final reference methylation atlas.
(4) CelFEER adapted CelFiE’s method with the following major modifications: (a) use 500 bp windows instead of CpG sites to find marker regions; (b) employ read averages (the ratio of methylated CpG sites to the total number of CpG sites on a read) instead of methylation ratio as the methylation quantification metric; (c) compare the level of each cell type to the minimum instead of the median methylation level of all other cell types; (d) select the top 150 markers before narrowing down to the top 100 markers for each cell type.
(5) UXM selected the top 25 specifically unmethylated genomic blocks with at least 5 CpGs using the *find_markers* function (--pval 0.05 -c 5 --min_bp 10 --max_bp 1500 --sort_by delta_means --tg_quant 0.25 --bg_quant 0.025 --top 25 --only_hypo) from wgbstools.

### Implementation of each cfDNA deconvolution method

To execute MethAtlas (https://github.com/nloyfer/meth_atlas), the *deconvolve.py* script was utilized, taking in silico cfDNA mixtures and reference methylation atlas as inputs for the deconvolution process. For cfNOMe (https://github.com/FlorianErger/cfNOMe), the *methylation_deconvolution.py* script was employed (--ineq --verbosefor --ref_header True – sample_header False) for deconvolution. For CelFiE (https://github.com/christacaggiano/celfie), *celfie.py* was used with --unknowns set to 0, unless explicitly specified. Similarly, for CelFEER (https://github.com/pi-zz-a/CelFEER), the *celfeer.py* was employed with --unknowns set to 0, unless otherwise specified. Lastly, for UXM (https://github.com/nloyfer/UXM_deconv), the *uxm deconv* command from UXM_deconv was executed for deconvolution.

### Evaluation of reference markers selection

To evaluate the cell type specificity and homeostasis of the reference markers selected by each deconvolution method, we downloaded cell-type-specific methylation marks (CpG sites) from MethyMark database (http://fame.edbc.org/methymark/) and DNA methylation variability data of immune cells from ImmuMethy (http://immudb.bjmu.edu.cn/immumethy/index.jsp). The MethyMark database integrates 50 methylomes across 42 human tissues/cell types, with 15 cell types overlapping with those in our reference methylation atlas (**Supplementary Table 5**). To quantify variability in ImmuMethy, we measured it in two ways: inter-quartile range (IQR) and standard deviations (SD). Four immune cell types (B cell, granulocyte, monocyte, and T cell) from ImmuMethy, having more than three conditions overlapping with the reference cell types in our study, were selected for subsequent analysis. SD and IQR across different conditions of the same cell types were calculated based on the mean methylation value of each condition.

For data from the MethyMark database, we first utilized liftOver from the UCSC Genome Browser to convert the genome coordinates of CpGs from hg19 to hg38. Subsequently, BEDTools^24^ (v2.31.0, bedtools intersect -wa -u) was employed to intersect the reference markers identified by each deconvolution method with cell-type-specific markers in MethyMark for each of the 15 overlapping cell types. The overlapped ratio was employed to assess the markers’ cell type specificity, with higher ratios indicating better specificity. Regarding data from the ImmuMethy database, BEDTools was also used to obtain SD and IQR values for the reference markers selected by each deconvolution method. Higher SD or IQR values corresponded to increased variability.

### Evaluation of sequencing depth filter thresholds in the reference methylation atlas

To evaluate the influence of sequencing depth filters, we implemented 13 depth filter thresholds (15, 20, 25, 30, 35, 40, 45, 50, 60, 70, 80, 90, and 100) during the generation of the reference methylation atlas. Specifically:

- For MethAtlas and cfNOMe, the reference methylation atlases were generated by applying each of the 13 thresholds to the median depth per CpG site across all samples.
- In case of CelFiE and CelFEER, reference methylation atlases were generated by adjusting the *depth_filter* parameter accordingly.
- For UXM, we simply modified the --min_cov parameter for the *find_markers* in wgbstools, keeping the other parameters unchanged.

### Evaluation of missing markers or cell types in the reference methylation atlas

To assess the impact of missing markers, we introduced five gradient missing proportions (10%, 20%, 30%, 40%, and 50%) for reference markers. This approach simulated scenarios where input information for deconvolution methods might be incomplete. The corresponding numbers of missing markers were randomly removed to represent a diverse range of genomic locations and ensure unbiased results. Each missing proportion was repeated 20 times to ensure the robustness of our findings. For the evaluation of missing cell types, we randomly deleted one cell type five times to generate five reference atlases with one missing cell type. Similarly, we deleted two cell types five times to generate five atlases with two missing ones. Subsequently, each deconvolution method was executed using reference methylation atlases with missing markers/cell types. All these analyses were done under a sequencing depth filter of 15.

### Assessment of running time and memory requirements

The running time and memory requirements were evaluated using memory-profiler (v0.61.0), a third-party Python package. For analysis, we selected a set of 100 cfDNA mixtures under uniform distribution with sequencing depth filter of 15 for the subsequent analysis. The total runtime and peak memory usage for all 100 samples were systematically recorded for each deconvolution method. To enhance the reliability of our results, we executed each deconvolution method ten times under consistent computational conditions: utilizing an Intel(R) Xeon(R) Gold 5218R processor with 40 threads and 192 GB of memory, operating on Ubuntu 22.04 LTS with Python version 3.10.11.

### Evaluation metrics

To assess the performance of each method, we computed both Pearson correlation coefficients and root mean squared errors (RMSE) between cell type proportions derived from each cfDNA mixture with known compositions. Higher Pearson correlation and lower RMSE values were indicative of superior deconvolution performance.

We used a scoring-ranking system based on two evaluation metrics (RMSE and Pearson correlation) to summarize benchmarking results across two datasets.

- *Accuracy*: For each dataset, we initially normalized the median values of RMSE and Pearson correlation (under a sequencing depth filter of 15) using the following equations:

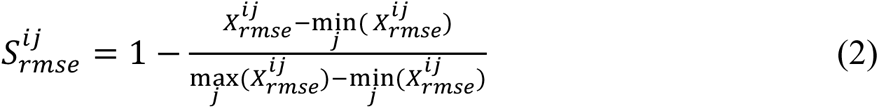

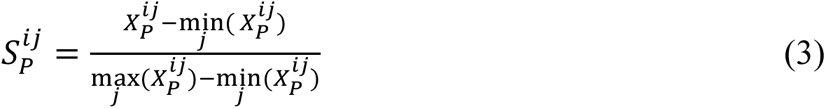

Here,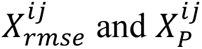 represent the median values of RMSE and Pearson correlation of method 𝑗 on the 𝑖-th dataset, respectively. Lower RMSE corresponds to higher values of 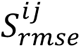, and higher Pearson correlation corresponds to higher values of 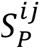. The scores were then aggregated across different evaluation metrics and datasets by calculating the arithmetic mean:

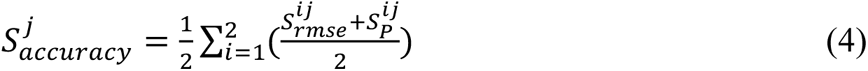
- *Robustness*: We first normalized the median values of RMSE and Pearson correlation for 100 mixtures to obtain a normalized score:

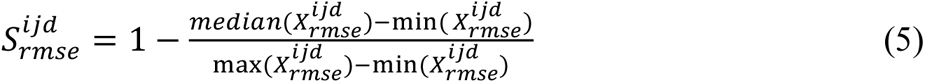

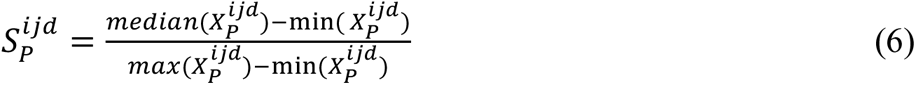

Here, 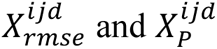 represent values of RMSE and Pearson correlation for method 𝑗 on dataset 𝑖 under evaluated factor 𝑑 (where 𝑑 represents 13 filter thresholds for *Depth filter*, six missing proportions for *Missing markers*, and 11 categories for *Missing cell types*, respectively). Then, the scores were aggregated across different evaluation metrics and datasets:

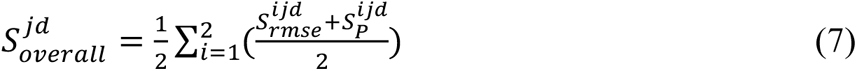 Subsequently, to assess the impact of *Depth filter* and *Missing markers* on the deconvolution results of each method, we calculated the variance of the above-defined 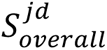 under different depth filter thresholds 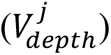 and missing markers proportions 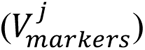, separately. For *Missing cell types*, the relative difference to the score with no cell type missing was calculated using:

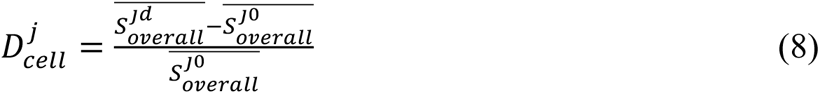

where 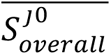 is the average score when there is no cell type missing, and 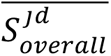 4is the average score for all others except no-cell-type missing category. A higher 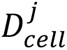 represents more robust performance.
- *Usability*: The aggregated score for running time 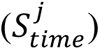 and memory requirements 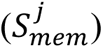 were calculated using the same approach as for *Accuracy*. Finally, we normalized the aggregated scores 𝑧^j^ of each evaluation factor for comparison and aggregation across different evaluation factors:

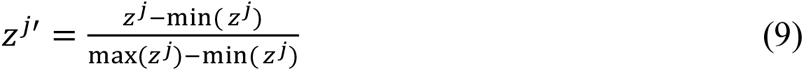 Here, 𝑧^j^ represents 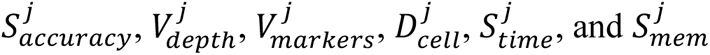 respectively. A higher (9) score corresponds to a better deconvolution performance. The overall performance for each method was calculated based on the normalized scores using 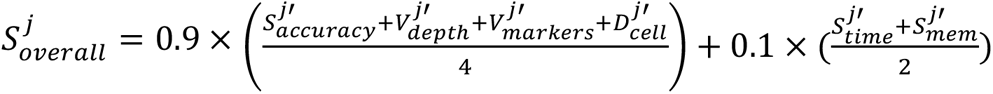. We assigned different weights because we think accuracy and robustness are much more important than usability for the users.

## Consent for publication

Not applicable.

## Author contributions

Y.L. conceived and supervised this project. T.S. performed most of the computational analysis.

J.Y. and Y.Z. performed part of the computational analyses. Y.L. wrote the manuscript. All authors interpreted the results, read, revised, and approved the final manuscript.

## Acknowledgements

The authors thank members of Yumei Li’s lab for helpful discussions. The authors also appreciated the insightful suggestions from members in Dr. Jingyi Jessica Li’s lab and and Dr. Wei Li’s lab.

## Supplementary information

Supplementary Figs. 1 to 7, and Tables 1 to 6.

