## Supplementary material for "Systematic Evaluation of Cell Type Deconvolution Methods for Plasma Cell-free DNA": Supplemantary figures 1-7

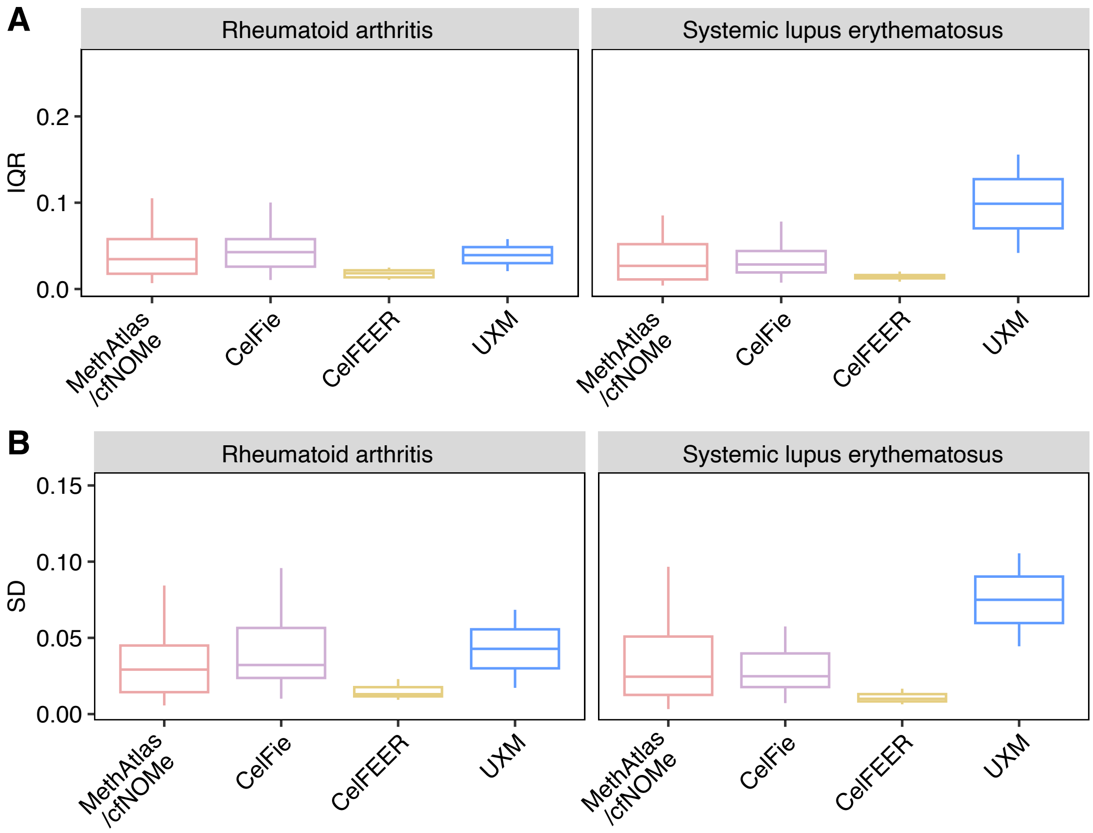


**Fig. S1.** The variability of the selected markers across individuals with B cell related diseases measured by Inter-Quartile Range (IQR) **(A)** and Standard Deviations (SD) **(B)**. IQR and SD data are sourced from the ImmuMethy database, focusing on B cell.

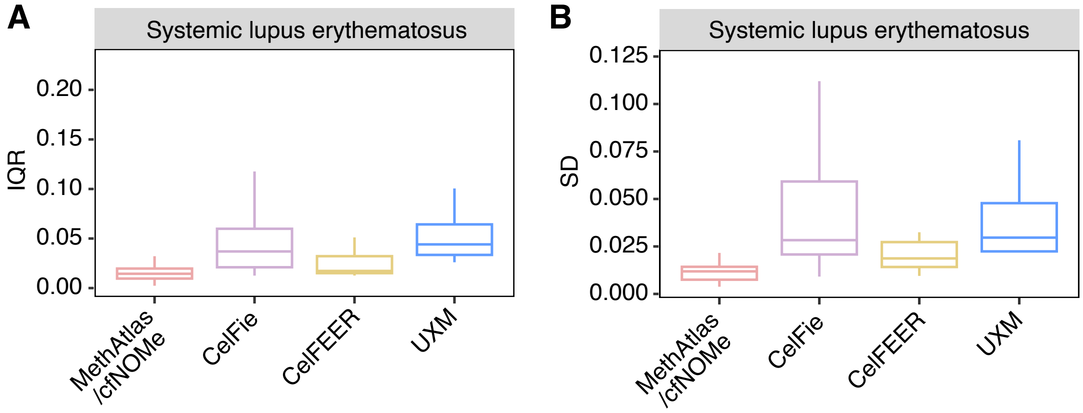


**Fig. S2.** The variability of the selected markers across individuals with Granulocyte related disease measured by Inter-Quartile Range (IQR) **(A)** and Standard Deviations (SD) **(B)**. IQR and SD data are sourced from the ImmuMethy database, focusing on Granulocyte.

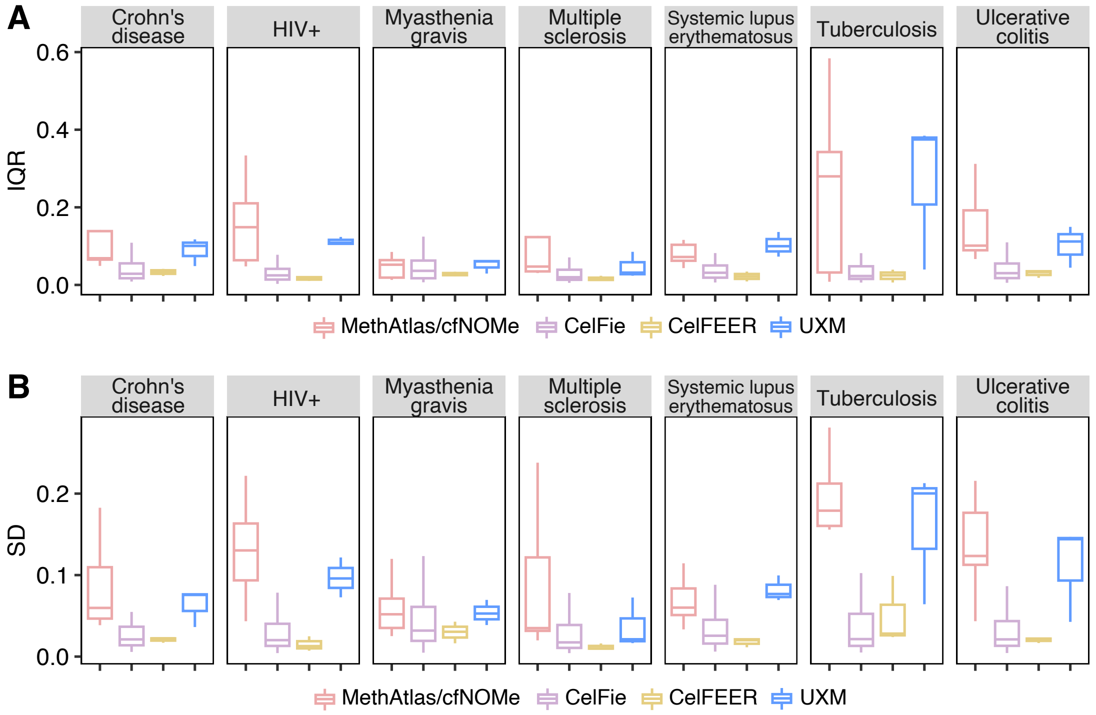


**Fig. S3.** The variability of the selected markers across individuals with Monocyte related diseases measured by Inter-Quartile Range (IQR) **(A)** and Standard Deviations (SD) **(B)**. IQR and SD data are sourced from the ImmuMethy database, focusing on Monocyte.

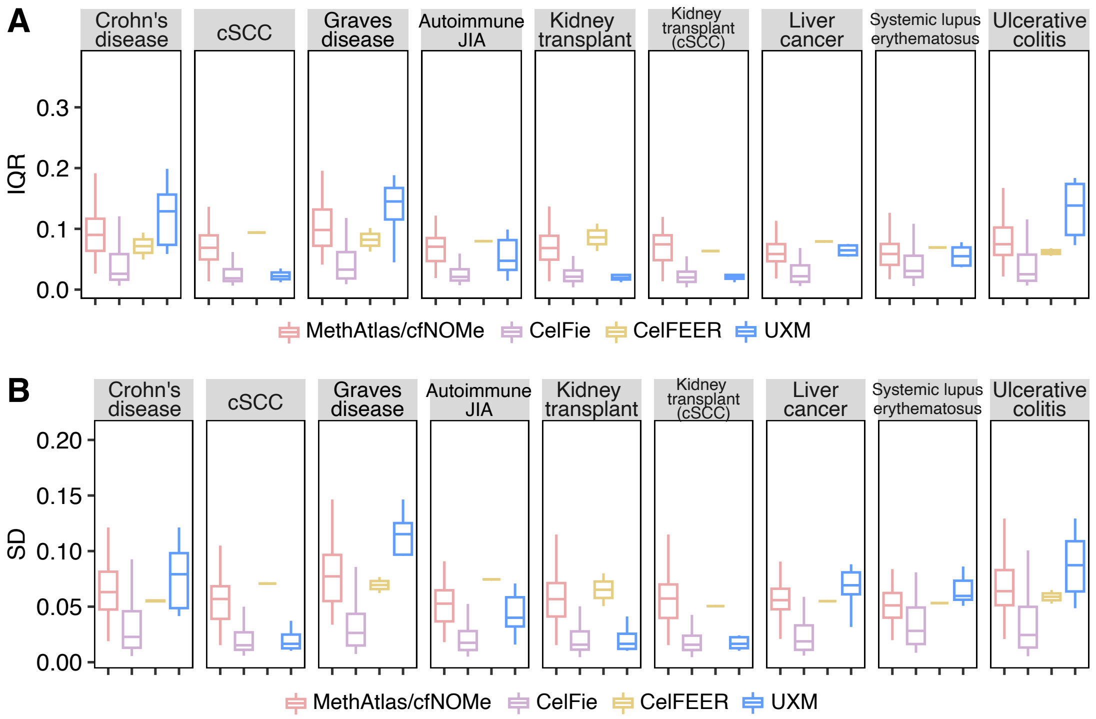


**Fig. S4.** The variability of the selected markers across individuals with T cell related diseases measured by Inter-Quartile Range (IQR) **(A)** and Standard Deviations (SD) **(B)**. IQR and SD data are sourced from the ImmuMethy database, focusing on T cell.

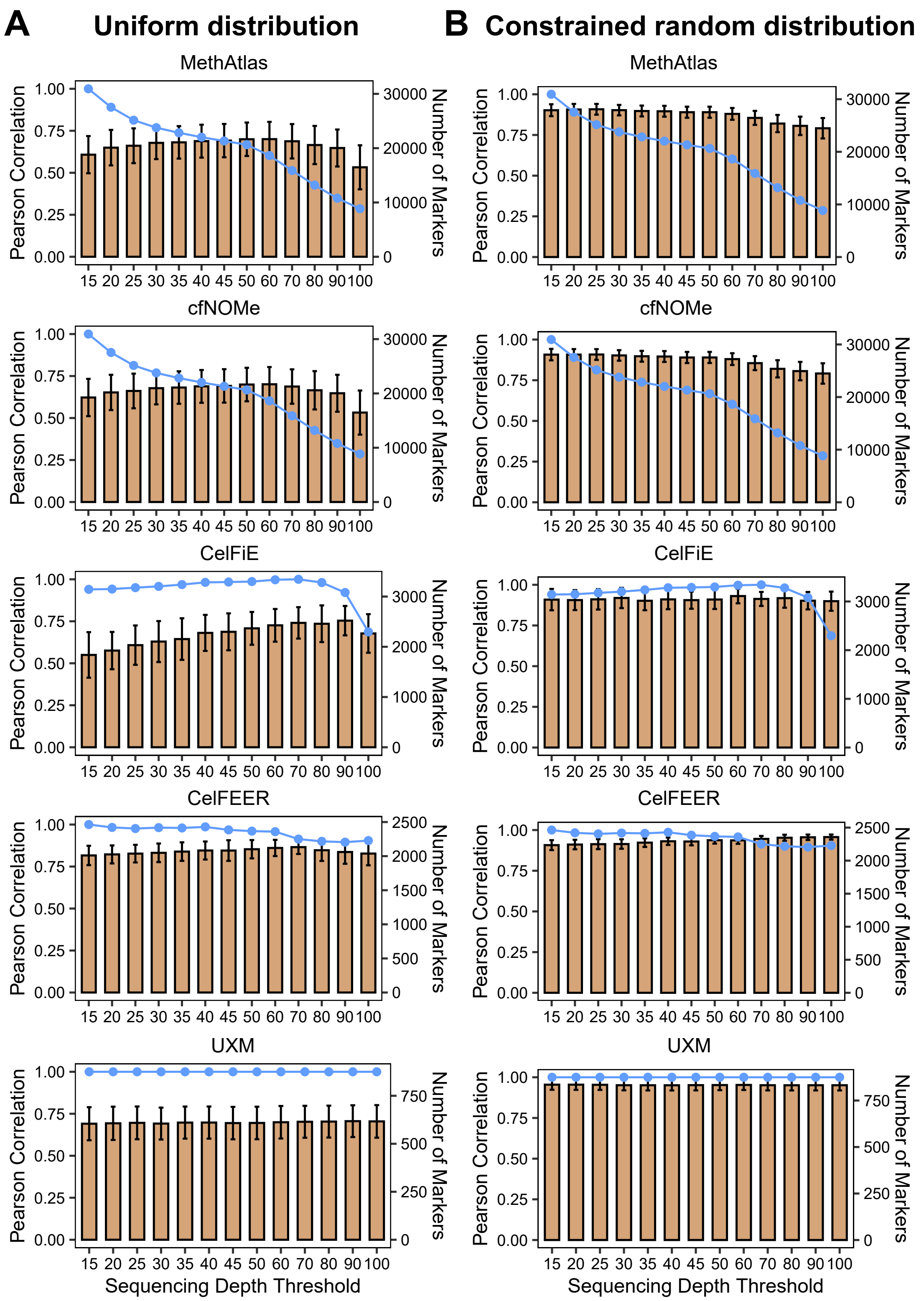


**Fig. S5.** Pearson correlations between the known proportions and the predicted proportions from different deconvolution methods, considering various sequencing depth thresholds for marker selection. The known proportions are generated under uniform **(A)** and constrained random distributions **(B)**. The line plot depicts the number of selected markers.

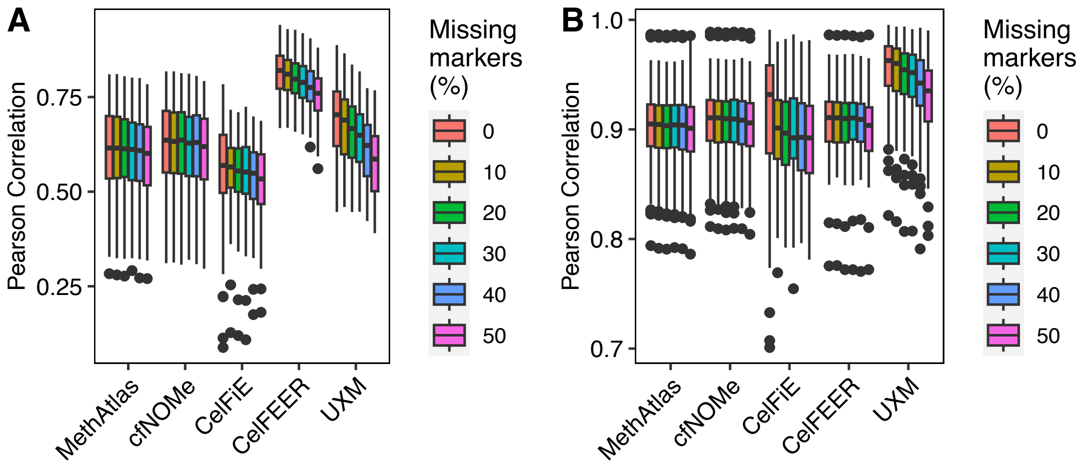


**Fig. S6.** Pearson correlations between the known proportions and the predicted proportions from various deconvolution methods. Different proportions of markers are intentionally omitted in the reference. The known proportions are generated under uniform **(A)** and constrained random distributions **(B)**.

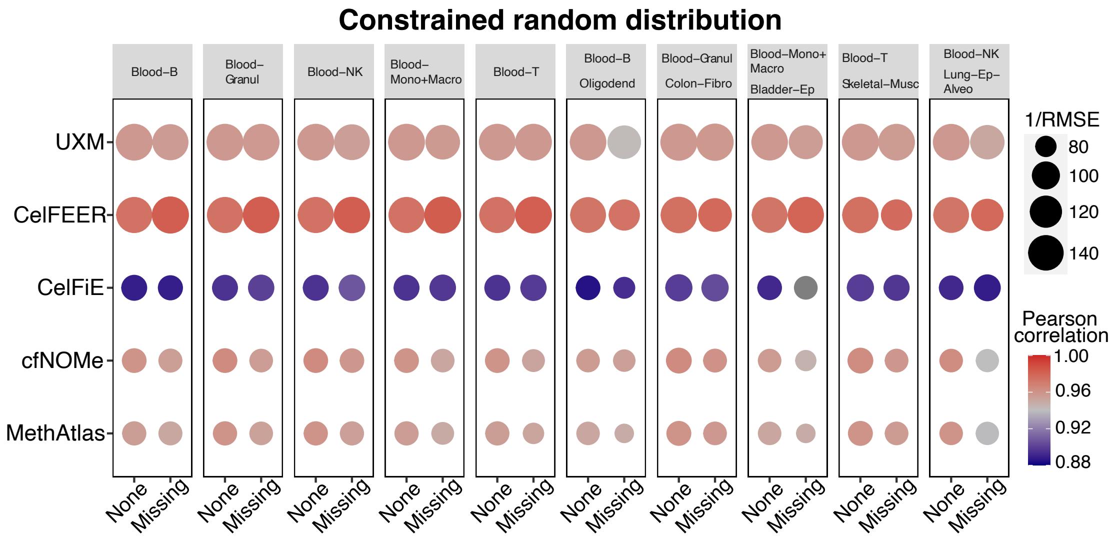


**Fig. S7.** Pearson correlations and RMSE values between the known proportions under constrained random distribution and the predicted proportions from different deconvolution methods with no, one, or two cell types intentionally missing in the reference.
